# The Taco Setup: A Novel TMS-fMRI Setup for High Resolution Whole Brain Imaging

**DOI:** 10.1101/2025.06.14.659622

**Authors:** Moataz Assem, Marius Mada, Steve Eldridge, Alexandra Woolgar

## Abstract

Simultaneous TMS-fMRI holds significant opportunities for advancing basic and translational neuroscience. However, current configurations face technical limitations, particularly the need to accommodate TMS hardware within the MRI environment. Previous solutions have used low numbers of radio-frequency (RF) channels, limiting fMRI data quality and whole-brain coverage. Here, we introduce a novel 22-channel “Taco” TMS-fMRI configuration that re-purposes flexible RF coils, wrapping them around both the participant’s head and the TMS coil to preserve whole brain signal reception. Guided by precision fMRI principles, we optimized acquisition protocols to achieve to achieve high temporal signal-to-noise ratio (tSNR) across the cortex. Data from three pilot participants demonstrate robust signal quality, including in regions proximal to the TMS coil. This setup offers a relatively simple and cost-effective approach to integrating precision fMRI into TMS-fMRI research.

## Introduction

Simultaneous transcranial magnetic stimulation and functional MRI (TMS-fMRI) is an emerging technique with strong potential for both basic neuroscience and translational applications—from causal mapping of brain networks to improving TMS-based diagnostics and interventions [1,2]. However, current TMS-fMRI setups face several technical limitations, particularly the spatial constraints imposed by fitting both the TMS coil and MRI radio-frequency (RF) receive coils close to the participant’s head. These constraints often necessitate reducing RF channel count, relative to gold standard fMRI, which in turn limits spatial coverage and signal quality.

Several solutions have been proposed, ranging from 1-channel birdcage coils that are large enough to accommodate the TMS coil and allow whole-brain imaging but with low data quality, to 7-channel surface coils that provide high signal-to-noise ratio (SNR) locally but inhomogenous coverage with relatively poor signal of brain regions distant from the coil [see [2,3] for comprehensive reviews]. New solutions are emerging (e.g., [4]) but so far the established approaches fall short of the SNR and coverage achievable with standard 32-channel head coils.

To address this gap, we developed a simple and cost-effective “Taco” TMS-fMRI setup (**Figure 1**). The design re-purposes two flexible body coils (18- and 4-channel), wrapped around the participant’s head and the TMS coil, yielding a 22-channel configuration. Guided by high quality fMRI acquisition recommendations [5], we optimized fMRI sequences on a 3T scanner to achieve whole-brain coverage with

**Figure 1.**
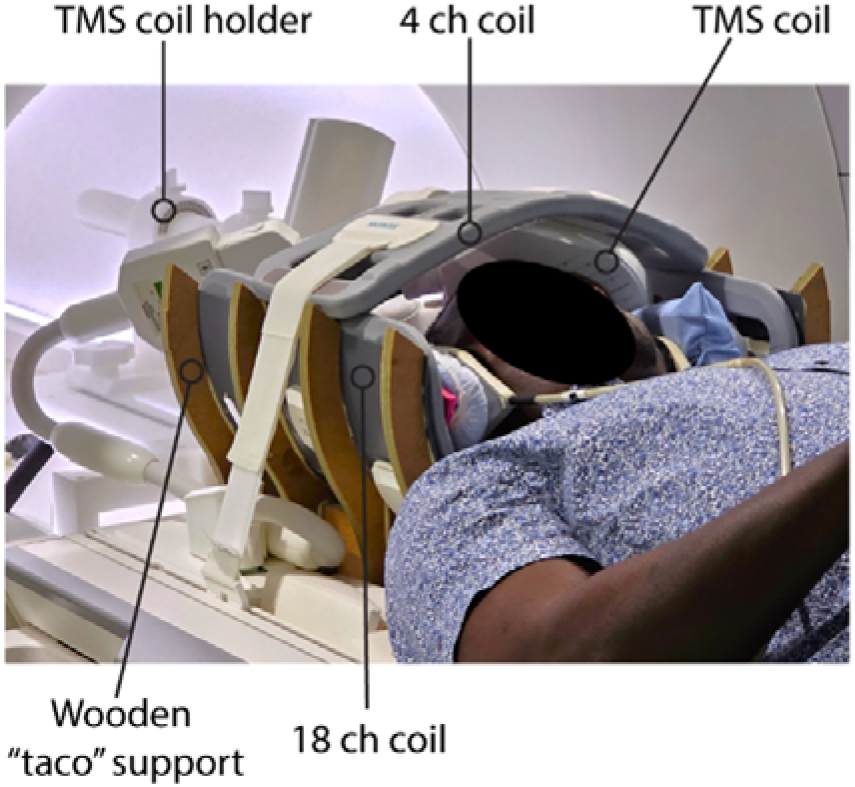
The “Taco” TMS-fMRI 22 channel setup. The configuration consists of two flexible RF coils—an 18-channel and a 4-channel coil—wrapped around the participant’s head and the TMS coil. The lower RF coil and the participant’s head rest within a custom wooden support shaped like a taco, which provides stability and consistent positioning across sessions. See **Methods** for additional details.

2.4 mm isotropic voxels, a TR of approximately 1 second, and a multiband acceleration factor of 4. In this initial report, we present pilot data from 3 participants showing high SNR across the cortex, including regions beneath the TMS coil. Additional data and analyses will be included as they are collected and processed.

## Methods

### Participants

We recruited 3 participants (3 males) who gave written informed consent, and data collection was approved by the University of Cambridge Ethics Committee (HBREC.2019.31).

### The Taco TMS-fMRI setup

The core design involved re-purposing two Siemens flexible RF coils—a 18-channel large flex coil and a 4-channel small flex coil—to form a closed 22 channel RF configuration around the participant’s head and the TMS coil.

To ensure consistent and stable coil placement across sessions, we constructed a custom prototype wooden headrest (see **Figure 1**). The large 18-channel RF coil was positioned on this base, and participants lay their heads directly onto it. For added comfort, a soft cushion was placed between the participant’s occiput and the RF coil.

The TMS coil (MagVenture MR-compatible air-cooled figure-of-eight coil, driven by a MagPro XP stimulator) was then placed directly over the left frontal scalp region. A ∼5 mm soft sponge pad was affixed to the contact surface of the TMS coil to enhance participant comfort. The TMS cable was routed through an RF waveguide into the scanner room and connected at the rear of the magnet bore. The coil was held in place using an MR-compatible MagVenture coil holder mounted inside the bore.

To complete the RF enclosure, the smaller 4-channel flex coil was carefully wrapped over the top of the participant’s head and the TMS coil, effectively “closing the taco.” Its edges were aligned with the borders of the lower large flex coil to form a continuous loop. The top coil was secured in place using Velcro straps attached to the MRI patient table. These straps also served to gently tighten and draw both RF coils closer to the participant’s head for maximal SNR during image acquisition. To minimize head motion during scanning, we placed foam cushions and head-stabilizing pads around the participant’s head within the setup.

Due to its compact size, the 4-channel coil allowed participants to view the projection screen via a standard MRI-compatible mirror mounted above the head, making the setup suitable for simultaneous TMS and task-based fMRI.

### Image Acquisition

All scans were acquired on a 3T Siemens Prisma scanner. Structural images were collected using a standard 32-channel RF receive head coil, following the Human Connectome Project (HCP) Lifespan imaging protocols [5–7]. These included one T1-weighted MPRAGE and one 3D T2-weighted SPACE scan, both acquired at 0.8 mm isotropic resolution.

For functional imaging, we were guided by the HCP 3T fMRI neuroimaging approach [5]. Our primary goals were to: (1) achieve whole-brain coverage (2) use voxel sizes equal to or smaller than 2.6 mm, approximately the mean cortical thickness, to reduce partial volume effects; (3) maintain a repetition time (TR) of equal to or less than 1000 ms to support better cleaning of fMRI artifacts.

The standard HCP 3T multiband echo-planar imaging (EPI) sequence uses 2-mm isotropic resolution, TR = 800 ms, TE = 37 ms, and a multiband (MB) acceleration factor of 8 [5–7]. However, this protocol was optimized for a 32-channel coil, whereas our setup involved a total of 22 channels.

We began by reducing the MB factor to 6, but found the resulting images unsatisfactory based on visual inspection. We subsequently tested MB4 at a range of voxel resolutions. For MB4, we tested 2.0 mm, 2.4 mm, and 2.6 mm isotropic resolutions (**Table 1**). Although we also evaluated MB2 sequences at multiple voxel resolutions, this lower acceleration did not meet our predefined criteria—namely, full brain coverage, voxel size ≤2.6 mm, and a TR ≤1000 ms and these data are therefore not included in this initial report.

**Table 1.** FMRI acquisition parameters tested with the Taco TMS-fMRI setup.

| Multiband factor | Number of slices | TR (ms) | Voxel size (mm) |
| --- | --- | --- | --- |
| MB4 | 48 | 1008 | 2.0 |
| MB4 | 52 | 1030 | 2.4 |
| MB4 | 48 | 1032 | 2.6 |

For all sequences, pre-scan normalize setting was set to “ON” to improve signal homogeneity across the brain. We also added a 100 ms gap at the end of each TR to simulate the post-volume acquisition delay typically used for artifact-free TMS pulse delivery (Note the TR values listed in **Table 1** do not include this additional 100 ms gap).

Each participant completed 2 resting-state fMRI (rfMRI) runs for each acquisition sequence. Each pair was acquired with reversed phase-encoding directions: one run in anterior-to-posterior (AP) and the other in posterior-to-anterior (PA) direction. For each sequence, we acquired 300 time points per run to allow direct comparisons between sequences. Additionally, spin-echo EPI images with matched geometry and reversed phase encoding (AP/PA) were acquired during both structural and functional sessions. The order of protocol testing was pseudo-randomized and counterbalanced across the three participants to minimize order effects.

### Data Preprocessing

The preprocessing followed the HCP minimal preprocessing pipelines (v4.0.0) [6] with some dataset-specific adjustments described previously [8,9] and briefly outlined below. Structural images (T1w and T2w) were used to extract cortical surfaces and segment subcortical structures, while functional images were mapped from volume to surface space and combined with subcortical data to form the standard CIFTI grayordinate space. Surface based spatial smoothing was applied using a 2-mm full-width at half-maximum (FWHM) kernel.

## Results

To provide an initial overview of signal quality, we computed temporal SNR (tSNR) maps for each acquisition (**Figure 2**). tSNR was calculated for each vertex by dividing the mean signal across the time series by its standard deviation. For every sequence, we first calculated tSNR maps for each individual run, then averaged the maps across the two runs per sequence. This was done separately for each acquisition condition.

**Figure 2.**
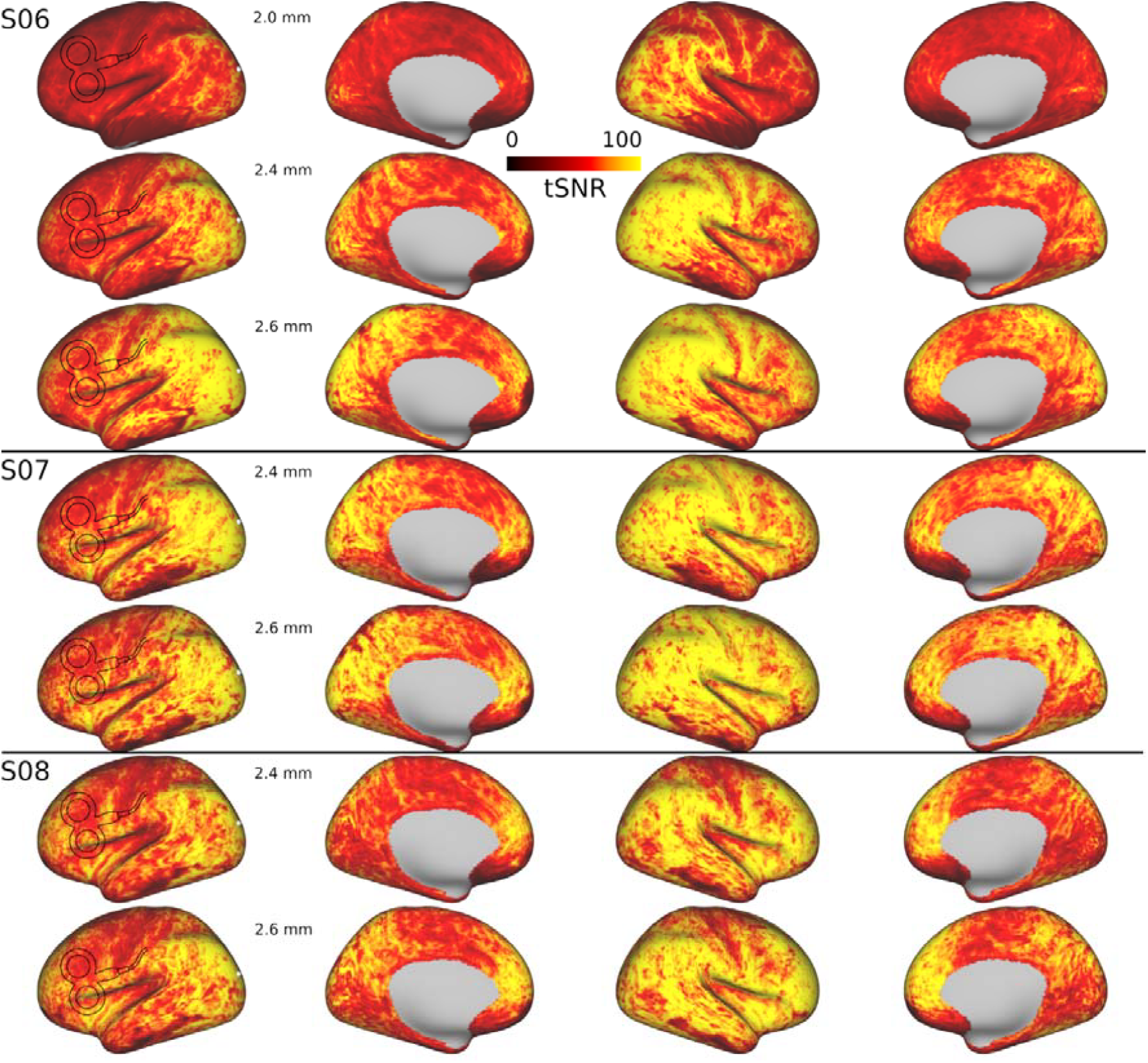
Resting-state tSNR maps for individual participants. Each column shows cortical tSNR maps for a different voxel resolution. The approximate position of the TMS coil over the left hemisphere is schematically illustrated in each map.

Figure 2. demonstrates that the Taco setup yields high tSNR values across the entire cortex. As expected, tSNR increased with larger voxel sizes—a trend evident in participant S06, where the 2.0 mm acquisition showed broadly lower tSNR compared to the 2.4 mm and 2.6 mm sequences. The difference between the 2.4 mm and 2.6 mm sequences, however, was minimal. A consistent hemispheric asymmetry was also observed, with lower tSNR in the left hemisphere, particularly in the frontal lobe, reflecting the placement of the TMS coil over that region. Despite this, tSNR values in the left frontal cortex remained within a robust double-digit range.

To quantify these differences, we extracted average tSNR values from several cortical areas. We used the HCP multimodal areal parcellation [9], which divides the cortex into 180 areas per hemisphere. We included primary visual area (V1), primary auditory area (A1), intraparietal sulcus (IP2), medial parietal cortex (POS2), fusiform face area (FFC), lateral prefrontal cortex (PFC) (p9-46v), medial PFC (8BM), and the insula (AVI). As shown in **Figure 3**, tSNR values for the 2.4 mm and 2.6 mm sequences were highly similar across all regions. However, both consistently showed reduced tSNR in the left hemisphere, particularly in areas directly beneath the TMS coil. Despite this reduction, tSNR in these regions remain high (≥ 50).

**Figure 3.**
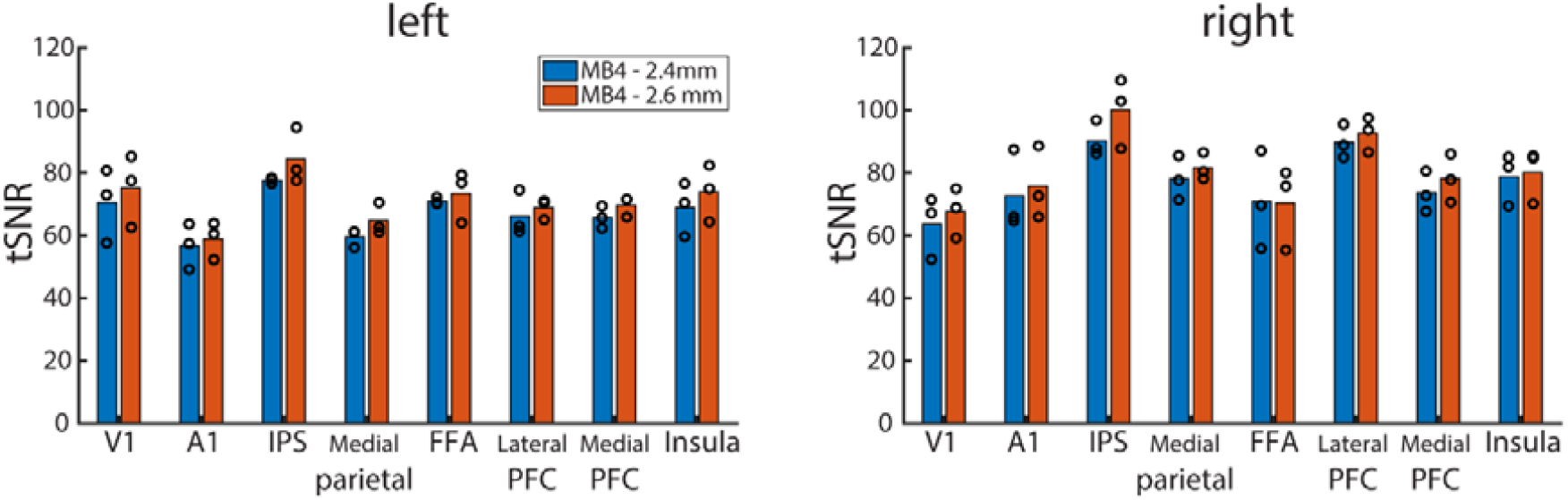
Areal tSNR. Each circle represents the tSNR value for one participant per cortical area.

## Discussion

This initial report demonstrates the feasibility and effectiveness of a novel, low-cost “Taco” TMS-fMRI setup that preserves whole-brain coverage and supports high-resolution fMRI. By re-purposing two flexible RF coils into a 22-channel configuration, we were able to implement precision fMRI acquisition protocols on a 3T scanner, achieving high tSNR values across the cortex—even in regions directly beneath the TMS coil. Compared to existing configurations [2], this setup offers a practical balance between hardware simplicity, participant comfort, and imaging quality. The data quality achieved was sufficient to support surface-based analyses, enabling more anatomically precise interpretations of TMS stimulation effects on fine-grained functional networks—an important step toward aligning TMS-fMRI with precision fMRI standards. While results are encouraging, ongoing work will evaluate setup performance across more participants, in comparison to other head coil configurations, during TMS stimulation and task-based paradigms. This report will be updated as additional data and analyses are completed.

## Acknowledgments

We would like to thank Matt Glasser and Marta Correia for useful discussions during the piloting phase. For the purpose of open access, the author has applied a Creative Commons Attribution (CC BY) license to any Author Accepted Manuscript version arising from this submission.

